# Blanking on Blanks: Few Insect Microbiota Studies Control for Contaminants - A Systematic Review

**DOI:** 10.1101/2024.08.30.610589

**Authors:** Elisabeth Williamson, Tobin Hammer, Katja Hogendoorn, Raphael Eisenhofer

## Abstract

Research on insect-microbe relationships is booming, with DNA sequencing being the most commonly used method to describe insect microbiota. However, sequencing is vulnerable to contamination, especially when the sample has low microbial biomass. Such low-biomass samples are common across insect taxa, developmental stages, and tissue types. Identifying putative contaminants is essential to distinguish between true microbiota and introduced contaminant DNA. It is therefore important that studies control for contamination, but how often this is done is unknown. To investigate the status quo of contamination control, we undertook a systematic literature review to quantify the prevalence of negative control usage and contamination control across the literature on insect microbiota (specifically bacterial communities) over the past 20 years. Two-thirds of the 243 insect microbiota studies evaluated did not include blanks (negative controls), and only 13.6% of the studies sequenced these blanks to control for contamination in their samples. Our findings highlight a major lack of contamination control in the field of insect microbiota research. This result suggests that a large number of microbes reported in the literature may be contaminants as opposed to insect-associated microbiota, and that more rigorous contamination control is needed to improve research reliability, validity and reproducibility. Based on our findings, we recommend a modified version of the guidelines outlined in the RIDES checklist: Report methodology, Include negative controls, Determine the level of contamination, Explore contamination downstream, and State the amount of off-target amplification.

**IMPORTANCE:** Our systematic review reveals a major lack of methodological rigour within the field of research on insect-associated microbiota. The small percentage of studies that control for contamination suggests that a considerable proportion of bacteria reported in the literature could be contaminants. The implication of this finding is that true microbiota may be masked or misrepresented, especially in insects with low microbial biomass.

## INTRODUCTION

Research on insect-associated microbial communities is booming globally. Communities of microbial symbionts (microbiota) can have profound ecological and evolutionary impacts on insects, that in some cases form obligate, species-specific symbiotic relationships with their host [e.g. 1, 2, 3]. However, most insect microbiota are yet to be described, or are in early stages of investigation, and the extent of microbial reliance and the nature of these associations across the insect world is unknown. The most commonly used method for characterising insect microbiota is DNA amplicon sequencing [4]. While DNA sequencing is a highly powerful, sensitive and accessible tool, interpretation of the data requires great care.

One of the biggest limitations for amplicon sequencing assessments is DNA contamination. Biological samples can become contaminated from exposure during collection, contact with research personnel and the laboratory environment, as well as from reagents, DNA extractions kits (called ‘kitomes’) and cross contamination between samples (called ‘splashome’) [5, 6, 7, 8]. When uncontrolled, DNA contamination can result in erroneous community assessments by distorting taxonomic diversity, obscuring differences between samples, and misrepresenting true absences of microbiota [9]. For example, human placental tissue was once thought to harbour specific microbiota, but later studies found that almost all of the sequence data could be attributed to contamination [6, 10, 11]. Contamination is particularly problematic when biological samples have a low number of microbes (called “low biomass samples”). Low biomass samples occur when microbes naturally exist in low abundance, or because the biomass of the sample itself is small. Low biomass samples are more prone to DNA contamination as it can represent a larger proportion of sequences in a given dataset [12, 5].

Universal primers designed to amplify sequences of bacterial 16S rRNA genes can also amplify chloroplast and mitochondrial genomes (referred to as ‘off-target DNA’) due to their common evolutionary history [13, 14]. Consequently, 16S sequencing assessments can experience amplification bias toward off-target DNA if samples are low biomass and contain animal or plant tissue. An abundance of off-target DNA can distort and obscure community assessments by reducing the representation of low abundance taxa, and this can further exacerbate the impact of contamination. Furthermore, there are several cases where high abundances of off-target DNA in insect samples have indicated low biomass samples, such as in *Lepidoptera* [15, 16], *Hymenoptera* [17], *Thysanoptera* [17], and *Phasmatodea* [18].

There are established protocols to control for DNA contamination. The standard approach, and the method focused on in this systematic review, is to process no-template or negative controls (i.e., blank samples) alongside biological samples during DNA extraction, PCR, and sequencing. Amplicon sequence variants (ASVs) can then be removed from the biological sample data if they are detected and more prevalent in negative controls [8, 12, 5]. The accuracy of such techniques can be further improved by employing statistical packages, such as Decontam [9], that classify ASVs as contaminants using established and reproducible methods.

Negative controls can also be used to measure the limit of detection (LoD). The LoD is a benchmark to determine the lowest amount of sample-derived DNA that can be reliably used to identify and quantify microbial taxa in a given dataset [19, 5]. It is measured using quantitative PCRs (qPCRs) to obtain the absolute abundance of negative controls which is then compared to the abundance from individual samples. Any sample that falls below the LoD can be discarded, and this process should be repeated per batch of samples as the degree of contamination can vary between lots processed [5, 12, 20].

Controlling for contamination is essential, but how often it is actually done in the insect microbiota literature is unknown. Here, we systematically assess whether insect microbiota research over the previous 20 years has appropriately controlled for DNA contamination. Specifically, we systematically address the following questions: 1) What percentage of studies have used negative controls to control contamination? 2) Has there been an increase in the proportion of studies that control for contamination over the years? 3) How many studies have determined their experimental limit of detection? 4) What proportion of studies acknowledge off-target amplification from chloroplast or mitochondrial DNA? In light of our findings, we provide recommendations to improve the robustness of future insect microbiota research.

## MATERIALS AND METHODS

### Search strategy, KAPPA analysis, and Filtering

This review was conducted according to the Preferred Reporting Items for Systematic Reviews and Meta-analyses Statement (PRISMA) [21]. We systematically searched Scopus, Web of Science and Google Scholar for relevant articles and data papers. Due to the lack of standardisation between electronic databases, we tailored our search strategy for each database to include certain terms and synonyms relating to our key concepts. Specifically, we searched for papers that described or investigated the microbiota of insects (searches provided in supplementary materials, Figure S1). Ordered by ‘relevance’, the first 200 search results from the three database searches were then imported into the Covidence platform [22] and duplicate papers were removed. Before filtering all imported studies using the Inclusion/Exclusion criteria (supplementary materials Table S2), 50 papers (sorted by ‘Title’) were assessed individually by two authors for comparison using a KAPPA analysis. Because the KAPPA analysis confirmed the screening was reproducible, we proceeded with the remaining articles to produce the final pool of papers for metadata collection. A summary of the information mined from our final pool is given in Figure 1, and the detailed data extraction method is given in the supplementary materials (Figures S3-S4).

**FIG 1.**
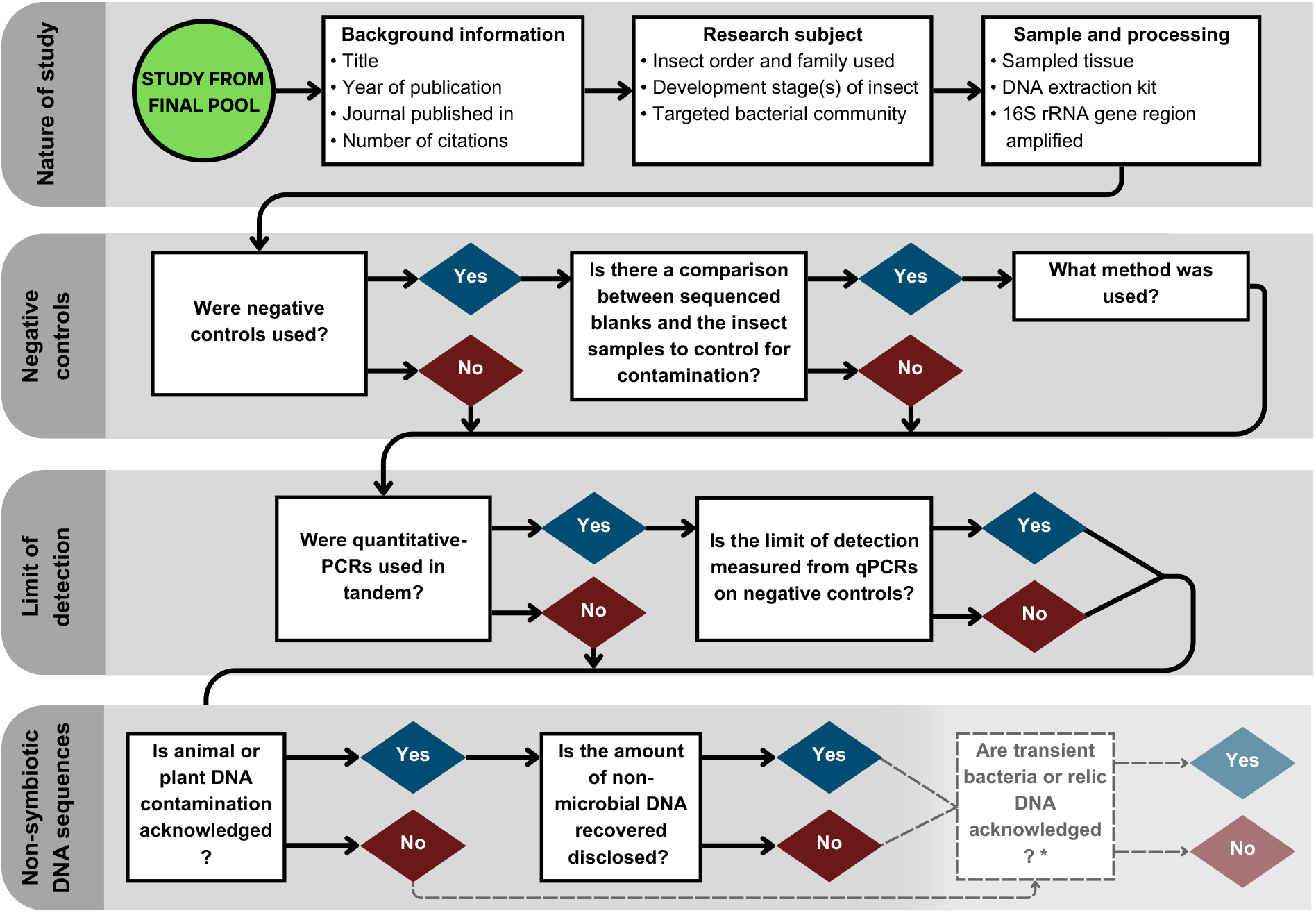
The information mined from the final pool of papers that researched insect microbiota using 16S sequencing to create the metadata set. *Additional questions addressed in the supplementary material.

### Metadata analysis

We first assessed the percentage of the most represented insect orders, to verify a lack of bias to certain taxa. We then examined trends in specimen sampling and processing by obtaining the percentages of the developmental stage(s) investigated (only adults, only juveniles, or multiple stages), the inclusion of a surface sterilisation step, and the region of the 16S rRNA gene used. It was determined how many studies mentioned mitochondria and/or chloroplasts in their methods or results sections, and if mentioned, whether they disclosed the amount of off-target amplification. We calculated the proportion of studies that reported using a negative control, and that used qPCRs to measure the limit of detection. From the subset of studies that used a negative control, we further calculated the proportion of studies that sequenced their controls and used this information to control for contamination. After assessing the distribution of citation counts using a Shapiro-Wilk test for normality, we opted to use a Mann-Whitney-Wilcoxon (MWM) test to assess whether the number of citations differed between studies that did or did not control for contamination, as an indicator of their perceived usefulness.

## RESULTS AND DISCUSSION

### Exploring the metadata

Our final pool consisted of 243 papers published between 2011-2022, which were used for metadata collection and subsequent analyses (supplementary materials Figure S5).

### Representation of insect orders and sampling approaches

The top 5 insect orders represented in our final pool were *Diptera* (23%), *Hemiptera* (18.1%), *Lepidoptera* (16.9%), *Coleoptera* (14.8%), and *Hymenoptera* (14%). The remaining 13.2% studied *Blattodea, Odonata, Orthoptera, Psocodea, Siphonaptera*, or multiple different insect orders. The distribution of insect orders studied indicates there was no bias to one field in our metadata. For the insects studied, 77.5% of studies sampled one metamorphic stage, with 51.4% of papers focusing on the adult stage, 25.9% sampling only juveniles (e.g. eggs, larvae, nymphs, or pupae), and the remaining 22.6% investigated more than one stage of development. A total of 56.8% of studies aimed to describe the microbial community of the gut specifically, 39.1% of studies described the community for the whole insect, and the remaining targeted specific organs or tissues.

### Trends in specimen processing methods

A total of 66.7% of studies included a surface sterilisation step. A range of substances were used to wash insects, with ethanol, bleach, sterile water, PBS, and detergent most frequently used. The variation in chemicals and whether or not insects were washed exemplifies the current lack in uniformity throughout insect-microbiota studies. In addition, many papers that included a surface sterilisation step stated that the purpose of the procedure was to remove external microbes. Contaminating microbes on the surface of specimens can be introduced during the handling and storage, or, occur naturally as a consequence of environmental contact (i.e. transient bacteria). It is possible that in some cases, surface sterilisation was presumed sufficient to control contamination. However, this fails to acknowledge downstream contaminants introduced during molecular processing, and previous research has shown no detectable effect of surface sterilisation on insect microbiota [23].

### Prevalence of negative control usage and controlling contamination in insect microbiota studies

Only one third (80/243) of studies reported using at least one negative control.

Negative controls were referred to as blanks, extraction blank controls (EBCs), no-template controls (NTCs), PCR blanks, sequencing controls, or surface washes (from insects or tool rinses). Fewer than half (33/80) of the studies that reported including a negative control subsequently sequenced these and compared the taxa found in their blanks to the corresponding insect samples. Therefore, only 13.6% of the studies included in the systematic review assessed their sequencing results for contaminating taxa. While the number of papers investigating insect microbiota has increased exponentially between 2011 and 2022, the proportion of studies that used a negative control or controlled contamination per year has not increased (Figure 2). Studies that did not control for contamination were not less frequently cited (p = 0.84), suggesting a lack of scrutiny when interpreting the conclusions derived from uncontrolled studies.

**FIG 2.**
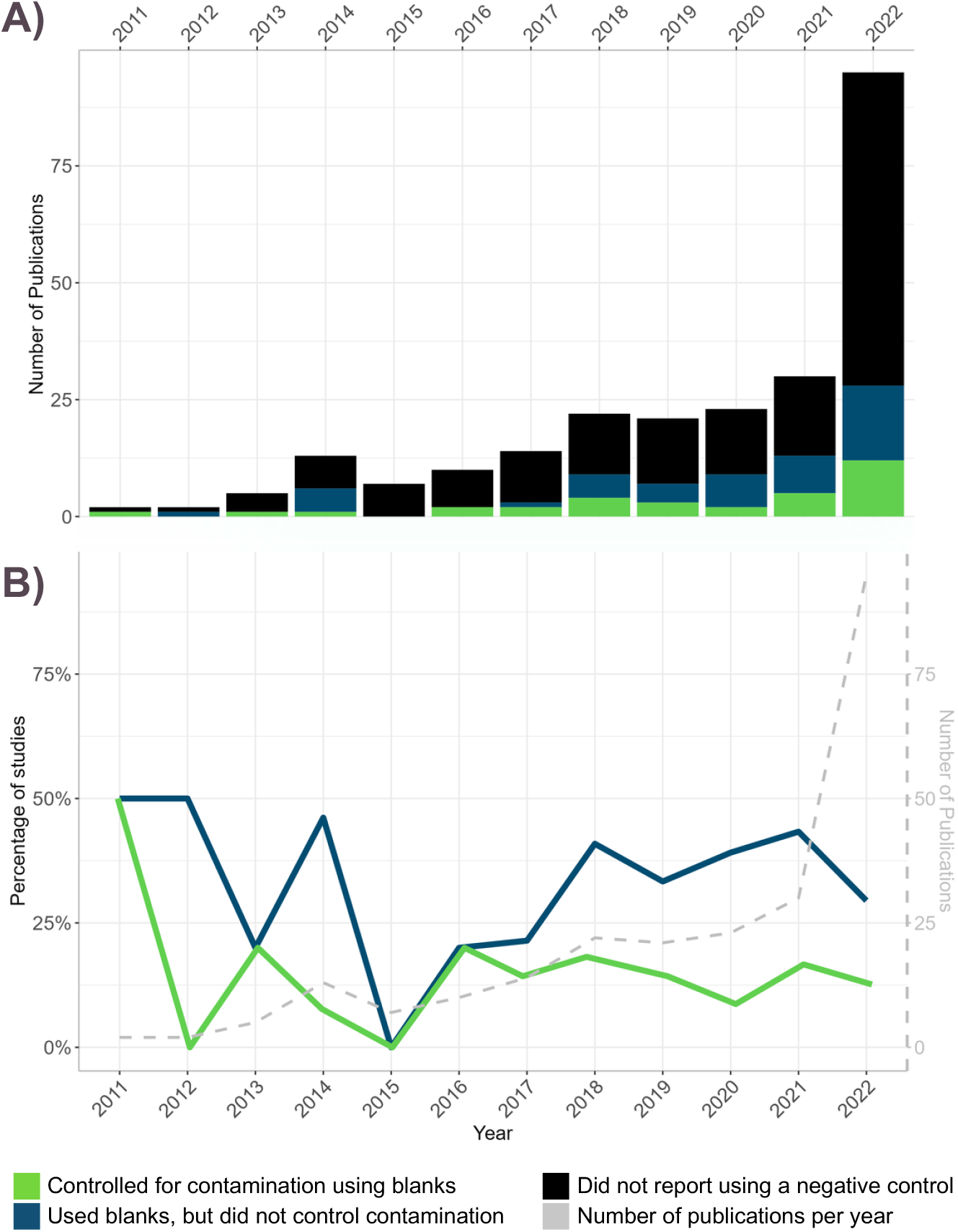
The trends in negative control use in our final pool of 243 from the past 20 years. A) The number of studies published per year that indicated using negative controls but did not control contamination (blue), used negative controls to control for contamination (green), and those that did not use negative controls (black). B) The percentage of studies per year that indicated using negative controls but did not control contamination (blue), that used negative controls to control for contamination (green), and the number of studies published (grey).

The low proportion of studies that report using negative controls (Figure 2) signifies that a large part of insect microbiota research has thus far not adhered to the latest guidelines that are recommended to control for contamination [5, 8, 24]. Although some studies may have used negative controls without mentioning them, this does not adhere to the latest recommendations and cannot be assumed.

The implication of this finding is that for 86.4% of studies included in our metadata, it can not be conclusively said whether the microbiota reported are truly insect-associated or the byproduct of contamination. One of the challenges in recognising contamination in insect studies is that some taxa of commonly reported insect microbiota overlap with that of common contaminants. For example, *Pseudomonas* [25], *Acinetobacter* [26, 27], *Staphylococcus* [28], *Bacillus* [29] and *Burkholderia* [30] are all described insect-associated microbes yet are also common contaminant taxa [12, 20, 5, 31].

Importantly, we also noted that there was considerable variation in the approaches used by the 33 studies that controlled for contamination. A total of 7/33 of studies used the statistical package Decontam [9] to call putative contaminant ASVs. The remaining studies used heuristic approaches, often involving a comparison between the communities identified, or manually filtering prevalent taxa found in negative controls from samples. Sometimes taxa were filtered based on the number of reads of putative contaminants from the controls, as a crude ‘limit of detection’ measure.

Without measuring the LoD using qPCR, it is impossible to reliably distinguish true taxa from the contamination-induced ‘background noise’ that is inevitably introduced during sampling and sequencing processes. Only 1.6% (n = 4 studies) measured the LoD using qPCR. The absence of measurements for the LoD, coupled with the few papers which controlled for contamination, further support the finding that few authors accounted for potential contamination in their results.

### Acknowledging off-target amplification from plant/animal DNA

Reporting the number of reads from off-target DNA is highly recommended to provide an accurate overview of the sequences recovered and to indicate the potential microbial biomass of samples and resulting sensitivity to introduced DNA contamination. We investigated what percentage of studies acknowledged chloroplast or mitochondrial DNA in their samples and found 35.2% included such acknowledgement. Most were studies that included a filtering step to remove chloroplast and/or mitochondrial DNA from their sequencing results without necessarily disclosing the extent of non-microbial DNA in the data, with only 5.8% (n = 14 studies) of total studies providing the amount of off-target amplification. It is possible that studies did not provide this information because off-target DNA represented few reads or was absent. However this is still informative, as disclosing the number of off-target DNA can also be useful when comparing 16S data between insect studies.

In addition to microbial biomass, the amount of off-target DNA amplification also depends on the diet of the specimen, gut content at time of sampling, and the primers used for PCR [13]. Therefore, there can be considerable variation in the amount of off-target DNA and amplification bias within insect species, depending on study design. The implication is that there can be differences in the bacterial abundances and diversity assessments, particularly in the representation of low abundance taxa, between studies. In these cases, reporting the proportion of off-target DNA can indicate the extent of the potential amplification bias and facilitate comparison between 16S datasets.

### Concluding remarks and recommendations

The vast majority of studies included in our systematic review did not follow best practice for controlling contamination. Consequently, a large proportion of published insect microbiota may be contaminated and this has the potential to mask and misrepresent true insect microbiota. While it is impossible to know the true number of studies affected, the lack of contamination control found in this systematic review suggests it could be high. This reveals a strong need for improved methodological rigour and adherence to standard guidelines across insect microbiota research. To address this, we recommend the RIDE checklist, a minimum standards guide for low microbial biomass studies that can be easily integrated into study design [5]. In addition to the four original guidelines, based on our findings we propose including an additional guide. The RIDES checklist stands for: 1) Report methodology, 2) Include controls in sequencing, 3) Determine the level of contamination using qPCRs to measure the limit of detection, 4) Explore the impacts of contamination in downstream analysis, and 5) State the amount of off-target DNA amplification to disclose potential amplification bias. The goals of these recommendations are to improve the accuracy and standardisation of future work, and reduce the uncertainty posed by undisclosed amplification bias and uncontrolled DNA contamination in the field of insect microbiota research.

## ACKNOWLEDGMENTS

R.E. conceived the project. R.E., T.H., and E.W., developed the methods. E.W. collected the metadata, performed the analysis, drafted the manuscript. E.W and R.E. designed the figures. E.W and R.E. performed the KAPPA analysis R.E and K.H supervised the statistical analyses. All authors discussed the results/implications and contributed toward the text of the final manuscript.

## DATA AVAILABILITY STATEMENT

All data on our filtering and metadata set are available in the supplementary materials. Code used to analyse the data is available on Github at: https://github.com/hi-its-lisou/Chapter4-BlanksSLR.git

## Supplemental material

- S1-S5 Search strategy and data extraction method
- SLR metadata, excluded papers, and KAPPA statistic
- Supplementary analysis: Additional considerations when assessing the nature of 16S sequences - Transient bacteria and Relic DNA

